# DAF-12 germline-to-soma signaling mediates transgenerational longevity in *C. elegans*

**DOI:** 10.64898/2026.08.05.743041

**Authors:** Scott P. Roques, Alexander K. Beaudoin, Jaime C. Croft, Tahreem Fiaz, Thiago Borges, Allison M. Sciarratta, Marybeth Slack, Teresa W. Lee

## Abstract

Development requires the complex coordination of gene regulatory networks that must remain robust in the face of variable environmental cues. In *Caenorhabditis elegans*, the nuclear hormone receptor DAF-12 integrates metabolic cues and hormonal signals to control important life history decisions, including development, reproduction, and the rate of aging. Here, we tested the involvement of DAF-12 germline-to-soma signaling in two transgenerational longevity mutants, *wdr-5* and *jhdm-1*. We have previously shown that both mutant populations gradually accumulate repressive H3K9me2 over multiple generations, which is necessary and sufficient for their lifespan extension. We find that *daf-12* activity was required for the epigenetic establishment of longevity in both mutant populations, but was only necessary for maintaining longevity in a *wdr-5* mutant background. Because DAF-12 also functions as a key regulator of dauer diapause, an alternative developmental stage triggered by environmental stress, we also tested the genetic relationship at earlier points in development. Surprisingly, mutations in either *wdr-5* or *jhdm-1* rescued the dauer defect of *daf-12* mutants, and we found a synergistic effect on unchallenged larval development in *wdr-5; daf-12* double mutants. These differing epistatic relationships indicate that, although the acquisition of longevity in both *wdr-5* and *jhdm-1* mutant populations shares a common mechanism, the impacts on somatic phenotypes (including lifespan extension) proceed via distinct pathways. Together, these results show how heritable chromatin states can co-opt existing developmental programs to influence key developmental decisions.

**ARTICLE SUMMARY:** How do early experiences influence development and aging? In this study, we explore this question by testing the genetic interaction between the DAF-12 signaling pathway and heritable chromatin landscapes. Previously, we showed that two *C. elegans* mutants can accumulate heterochromatin over multiple generations to acquire longevity. We find that DAF-12 is required to establish this epigenetic trait but is not necessary to maintain it. We also find that chromatin landscapes bypass DAF-12’s role earlier in development, including during the decision to enter dauer diapause. Overall, this study shows how chromatin states co-opt existing developmental programs to influence key life history decisions.

## INTRODUCTION

Organismal development must progress through stereotyped stages despite fluctuations in the environment. Therefore, a series of highly coordinated gene regulatory networks have evolved to maintain this robustness in the face of variable inputs. However, the same genetic programs that protect against adverse conditions early in life can become maladaptive if they persist into adulthood. The recently described programmatic theory of aging explores this concept further, unifying several prior hypotheses to explain how early developmental events act as the origins of both aging and adult diseases (Gluckman and Hanson 2004; De Magalhães and Church 2005; Blagosklonny 2006; Langley-Evans 2006; Gems 2022). Across animal taxa, changes in developmental environments (like larval or fetal exposure to nutrition) can impact adult health by causing changes in metabolism, growth, cognition, or reproduction (Godfrey et al. 2007). In the nematode *Caenorhabditis elegans*, environmental cues trigger larval arrest, hasten growth, or even dictate investment in the germline (Hibshman et al. 2016; Petrascheck and Miller 2017; Baugh and Hu 2020; Rashid et al. 2020). Some early developmental pathways are shared with those used later in life to promote healthy aging or defend against stress (Silva-García 2023) – two of the most well-studied aging pathways are also important for organismal development: insulin-like signaling and mTOR signaling. Because both pathways act downstream of nutrient sensing, lifespan and development are sensitive to changes in diet and the environment (Kaletsky and Murphy 2010; Kenyon 2010; Fernandes and Demetriades 2021).

Chromatin modifying enzymes play an important role in mediating transcriptomic changes during life transitions, including those that occur during development or aging (Allis and Jenuwein 2016); the reversible nature of chromatin modifications allows the epigenome to adapt to changing environmental cues in real time (Cui and Han 2007; González-Aguilera et al. 2014). For example, changes in chromatin state help to mediate the effects of nutrition on lifespan, while also impacting metabolic regulation in turn (Han et al. 2017; Johnson and Stolzing 2019; Silva-García et al. 2023; Emerson et al. 2024). Chromatin modifications are a cause and a consequence of gene expression: in some contexts, they are able to establish new transcriptional programs, whereas in others, they only serve as a reflection of existing transcriptional states. The methylation of histone H3 at lysine 4 (H3K4me) promotes gene expression and is also deposited co-transcriptionally by the MLL/COMPASS complex (Wood et al. 2007; Bannister and Kouzarides 2011; Henikoff and Shilatifard 2011; González-Aguilera et al. 2014; Benayoun et al. 2015; Soares et al. 2017; Millán-Zambrano et al. 2022). One essential component of MLL/COMPASS is WDR-5, which is conserved from yeast to humans (Dou et al. 2006; Steward et al. 2006); animals that lack *wdr-5* activity experience developmental defects, delay, or lethality (Byrd and Shearn 2003; Wysocka et al. 2005; Simonet et al. 2007; Vilhais-Neto et al. 2017). In contrast, methylation on histone H3 of the nearby lysine 9 (H3K9me) is associated with gene repression, with H3K9me2 most closely associated with canonical heterochromatin factors in *C. elegans* (Liu et al. 2011; Garrigues et al. 2015; Mcmurchy et al. 2017).

With each new generation, the vast majority of chromatin marks are removed during the process of epigenetic reprogramming, which provides embryos with an epigenetic reset that bestows totipotency for successful development (Kawashima and Berger 2014; Nashun et al. 2015). Histone modifying enzymes are necessary during this process to properly reestablish a ground state of active H3K4me and repressive H3K9me. The failure to properly reprogram these histone modifications has severe consequences: in mice, embryos lacking the H3K4 demethylase LSD1/KDM1A or H3K9 methyltransferase SETDB1 die early in embryogenesis (Dodge et al. 2004; Ancelin et al. 2016; Wasson et al. 2016; Zeng et al. 2026); in *C. elegans*, mutants suffer sterility and a severe developmental delay caused by the inappropriate expression of germline genes in somatic cells(Katz et al. 2009; Greer et al. 2014; Kerr et al. 2014; Carpenter et al. 2021).

We have previously shown that animals lacking the MLL/COMPASS complex gradually accumulate repressive H3K9me2 over many generations, likely due to the continued absence of active H3K4me during epigenetic reprogramming (Lee et al. 2019). This transgenerational accumulation of H3K9me2 is both necessary and sufficient for longevity. H3K9me2 accumulation also confers longevity in *jhdm-1* mutants, which lack a putative H3K9 demethylase orthologous to *S. pombe* Epe1 (Ragunathan et al. 2015; Lee et al. 2019). Notably, the transgenerational dynamics of longevity differs between these mutant backgrounds: *wdr-5* mutant populations become long-lived after twenty or more generations, whereas *jhdm-1* mutant populations are usually long-lived after six to eight generations, perhaps because JHDM-1 has a more direct effect on repressive H3K9me (Lee et al. 2019; Croft et al. 2023).

Intriguingly, after longevity has appeared in either *wdr-5* or *jhdm-1* mutant populations, it is heritable for four generations, even by descendants that are genetically wild-type (Greer et al. 2011; Lee et al. 2019). We propose a model in which accumulated heterochromatin can be inherited by descendants, although a balance of H3K4me and H3K9me is regained after four rounds of epigenetic reprogramming under wild-type conditions. However, it is still not clear why the inheritance of a repressive chromatin landscape affects lifespan – how can heterochromatin that accumulates in the germline influence aging, which mostly occurs in somatic tissues?

Developmental timing is tightly regulated by heterochronic genes, some of which are conserved across animal taxa (Ambros and Horvitz 1984; Ambros 1989; Lee et al. 1993; Wightman et al. 1993; Pasquinelli et al. 2000; Moss 2007). A major regulator of heterochrony in *C. elegans* is the highly conserved nuclear hormone receptor DAF-12, which, in vertebrates, is most homologous to two receptors involved in metabolic regulation: the vitamin D receptor and the liver X receptor (Antebi et al. 2000). DAF-12 sits at the a nexus of both metabolism and development, integrating environmental cues with hormonal signals to control growth and reproductive development (Riddle and Albert 1997; Antebi et al. 2000; Mooijaart et al. 2005; Motola et al. 2006; Hochbaum et al. 2011). If *C. elegans* experience unfavorable conditions as L1 larvae (like food depletion, crowding, or elevated temperature), they enter a developmentally arrested L3 larval stage called dauer (Cassada and Russell 1975; Riddle et al. 1981).

Dauer entry is facilitated by pheromones that reduce signaling in the insulin-like/DAF-2 and TGF-β/DAF-7 pathways, allowing DAF-12 to activate dauer genes and repress those needed for reproductive adulthood (Ren et al. 1996; Ogg et al. 1997; Gems et al. 1998; Antebi 2013). Dauer larvae undergo striking morphological changes (including a thickened cuticle and sealing of orifices) and exhibit specialized behaviors (like nictation or reduced locomotion) (Cassada and Russell 1975; Albert and Riddle 1983). Once a dauer individual encounters more favorable conditions, it resumes development to become a reproductive adult.

Dauer entry is adaptive because it allows for the preservation of fertility, although effects on reproduction differ depending on how dauer was induced (Hirsh et al. 1976; Hall et al. 2010; Ow et al. 2018). As a tissue, the germline is especially attuned to nutrient availability (Seidel and Kimble 2011; Laws and Drummond-Barbosa 2017; Galan et al. 2020; Webster et al. 2022). For example, in *C. elegans*, maintenance of the germline stem cell niche is closely tied to nutritional availability before dauer entry (Koitz et al. 2026). In *Drosophila*, oocyte differentiation is regulated (in part through the activity of another nuclear hormone receptor complex) (Finger et al. 2021; Bradshaw et al. 2024; Jung et al. 2026; Zike et al. 2026). Later in life, DAF-12 is involved in germline-to-soma communication that confers lifespan extension (Antebi 2013), including in *spr-5* mutants, which lack a H3K4 demethylase that has homology to mammalian KDM1A/LSD1 (Katz et al. 2009; Greer et al. 2016).

In this study, we tested whether DAF-12 germline signaling is involved in the transgenerational lifespan extension of *wdr-5* and *jhdm-1* mutants. We found that DAF-12 is necessary for establishing longevity in both mutant backgrounds but only required to maintain longevity in *wdr-5* mutants. Similarly, DAF-12 enhanced a developmental delay in *wdr-5* mutants, but not in *jhdm-1* mutants. Together, these findings suggested how differences in chromatin landscapes between these genetic backgrounds may affect germline-to-soma signaling. Surprisingly, we found that mutations in either *wdr-5* or *jhdm-1* could rescue the dauer-deficient phenotype of *daf-12* mutants, highlighting how chromatin landscapes impact DAF-12 function at important life history decisions. We show how gene programs used during early development may be affected by heritable chromatin states to affect an individual’s biology throughout its life cycle.

## METHODS

### Strains and husbandry

All strains were cultured using standard methods (Brenner 1974) at 20°C on 6-cm nematode growth media (NGM) plates seeded with OP50 *E. coli* grown overnight at 37°C in Luria Broth (LB). The following strains were obtained from the *Caenorhabditis* Genetics Center (RRID:SCR_007341), with sequence information provided by WormBase (Sternberg et al. 2024) and the Alliance of Genome Resources (Aleksander et al. 2024).

- N2: wild type (Bristol)
- RB1826: *jhdm-1 (ok2364) III*
- RB1304: *wdr-5 (ok1417) III*
- DR20: *daf-12 (m20) X*
- AA86: *daf-12 (rh61rh411) X*
- TER1: *jhdm-1 (ok2364) III; daf-12 (m20) X*
- TER2: *wdr-5 (ok1417) III; daf-12 (m20) X*
- TER20: *jhdm-1 (ok2364) III; daf-12 (rh61rh411) X*

### Transgenerational experiments

For each strain, three L4 hermaphrodites were transferred every third day from the previous population, except for *wdr-5* populations, in which four to six gravid young adults were transferred to ensure the selection of fertile animals. The P0 generation was the progeny of L1 animals recovered after a thaw.

### Lifespan

Assays were performed at 20°C on NGM agar plates that did not contain 5-fluoro-2’-deoxyuridine (FUdR). On Day 1, young adults (on their first day of egg-laying) were allowed to lay for 4–6 hours to hatch a synchronized population for the assay. When progeny were L4s or young adults, 90 animals per condition were transferred to new plates, with 30 animals per plate. Animals were transferred every day or every other day during their fertile period (usually the first ten days of adulthood). Plates were scored daily and animals marked as dead if they did not move in response to repeated prodding with a platinum pick. Animals were censored from analysis if they died from ruptured vulvas, matricide (“bag of worms” phenotype), or crawling off the agar. Kaplan-Meier survival curves were generated in GraphPad Prism and significance was calculated comparing generation-matched populations using a log rank test (Mantel-Cox). Lifespan differences are reported as percentages of median lifespans. The following key observations were repeated in blinded experiments: the effect of a *daf-12* mutation on establishment of longevity (Fig. 1), the effect of a *daf-12* mutation on the maintenance of longevity (Fig. 2), and the acquisition of transgenerational longevity (Supplementary Fig. 1).

**Figure 1.**
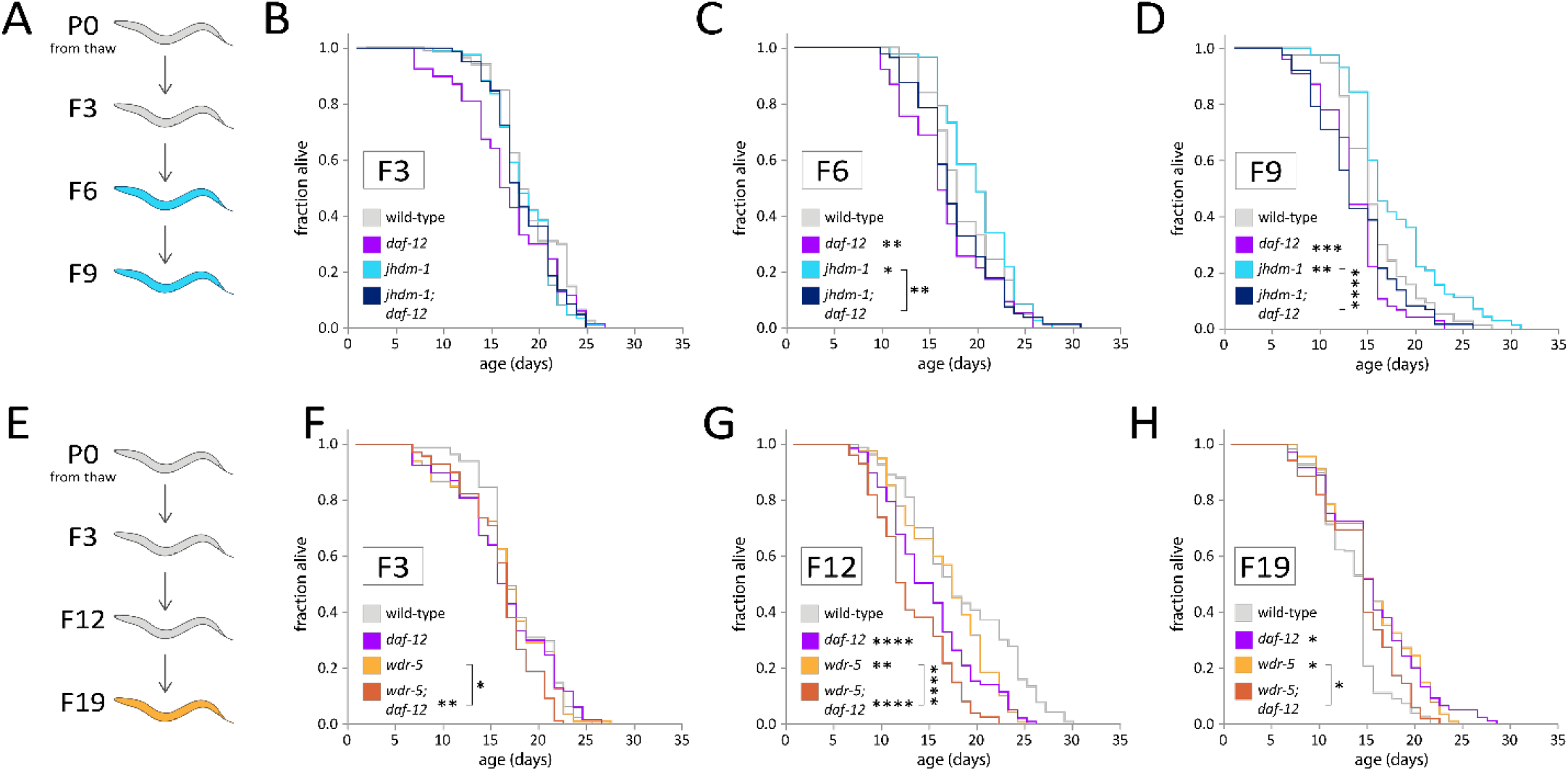
Establishment of longevity in *jhdm-1* and *wdr-5* mutants requires *daf-12* activity. (**A, E**) Schematic of transgenerational populations, with color of animals indicating acquisition of longevity in *jhdm-1* mutants (**A**, light blue) or *wdr-5* mutants (**E**, yellow). (**B-D**) Lifespan of F3 (**B**), F6 (**C**), or F9 (**D**) *jhdm-1; daf-12* double mutants (dark blue) compared to wild-type (gray), *daf-12* mutants (purple), or *jhdm-1* mutants (light blue). (**F-H**), F3 (**F**), F12 (**G**), and F19 (**H**) *wdr-5; daf-12* double mutants (dark orange) compared to wild-type (gray), *daf-12* mutants (purple), or *wdr-5* mutants (yellow). Statistical analysis was performed using a log-rank test compared to generation-matched wild-type populations (shown next to genotype) or between genotypes (indicated by bracket) (\**P* < 0.05, ** *P* < 0.01, \*\*\**P* < 0.001, \*\*\*\**P* < 0.0001). Median lifespan, statistics, and additional replicates are shown in Supplementary Table S1.

**Figure 2.**
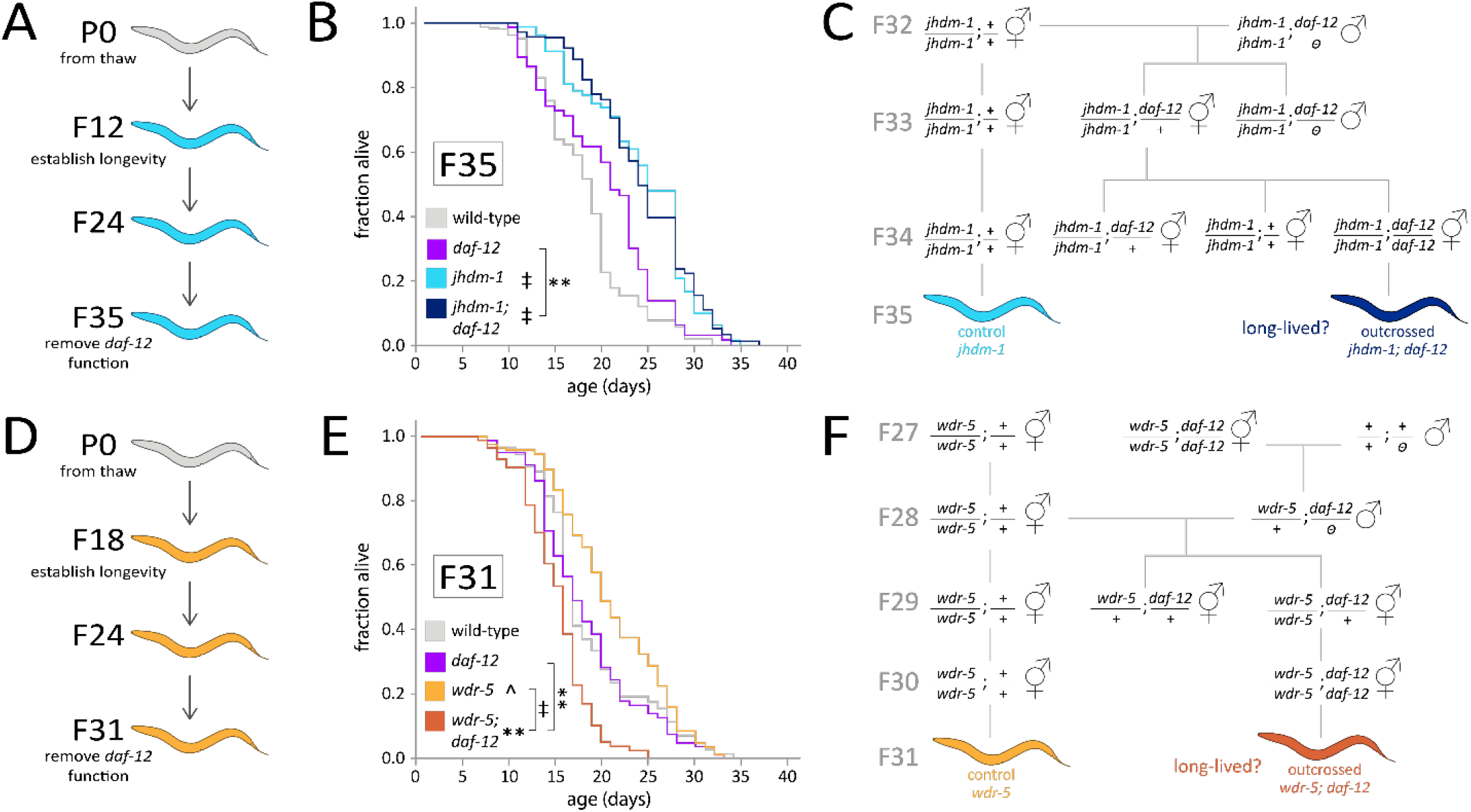
*daf-12* activity is necessary to maintain longevity in *wdr-5* mutants but not in *jhdm-1* mutants. (**A, D**) Schematic of transgenerational populations, with color of animals indicating acquisition of longevity in *jhdm-1* mutants (**A**, light blue) or *wdr-5* mutants (**E**, yellow); *daf-12* function was genetically removed in the generation indicated. (**B**) Lifespan of *jhdm-1; daf-12* double mutants (dark blue) in F35, a population in which *jhdm-1* single mutants (light blue) have acquired longevity, compared to wild-type (gray) and *daf-12* mutants (purple). (**C**, **F**) Genetic schemes for generating late-gen double mutants lacking *daf-12* activity (and the appropriate single mutant control) in a *jhdm-1* mutant background (**C**) or *wdr-5* mutant background **(F**). (**E**) Lifespan of *wdr-5; daf-12* double mutants (dark orange) in F31, a population in which *wdr-5* single mutants (yellow) have acquired longevity, compared to wild-type (gray) and *daf-12* mutants (purple). Statistical analysis was performed using a log-rank test compared to generation-matched wild-type populations (shown next to genotype) or between genotypes as indicated by bracket (^*P* = 0.05, \*\**P* < 0.01, ‡*P* < 0.0001). Median lifespan, statistics, and additional replicates are shown in Supplementary Table S2.

### Dauer entry

Synchronized populations were generated by allowing 5-10 young adult hermaphrodites to lay for 3-5 hours, after which mothers were removed from plates. Plates were kept at 27°C for 44 hours (Antebi et al. 2000). Populations were visually scored for dauer stage, as well as for L1 arrest (not reported), based on body width, body length, and dauer-like behavior. Each biological replicate included 3-12 technical replicates (individual plates of synchronized populations); 9 biological replicates were collected for *jhdm-1* mutant experiments, and 4 biological replicates were collected for *wdr-5* mutant experiments. Significance was calculated using an ANOVA comparing biological replicates with a post-hoc Tukey test. To verify the presence of dauer animals in the population, plates were washed with 1 mL of 1% SDS and gently agitated for 30 minutes at room temperature. Animals that were still alive and thrashing were considered dauer.

### Progeny counts

Individual hermaphrodites were cloned as L4s and transferred daily until no longer fertile. Progeny were scored as L4s or young adults. Each biological replicated started with examining broods from at least 5 mothers, although broods were censored if mothers died before the end of their laying period; 9 biological replicates were completed. Significance was calculated using an ANOVA comparing biological replicates with a post-hoc Tukey test.

### Larval development

Synchronized populations were generated by allowing 5 young adult hermaphrodites to lay for 3-5 hours, after which mothers were removed from plates. Plates were scored every 24 hours for three days; all animals were counted, then scored for larval stage. At least 5 technical replicates were included in each biological replicate (individual plates of synchronized populations); 4 biological replicates were collected for *jhdm-1* mutant experiments, and 3 biological replicates were collected for *wdr-5* mutant experiments. Significance was calculated using a Fisher’s exact test comparing age-matched populations.

## RESULTS

### The establishment of transgenerational longevity requires *daf-12* activity

We have previously shown that populations that accumulate heterochromatin over multiple generations gradually become long-lived (Lee et al. 2019; Croft et al. 2023). For example, mutants lacking the putative H3K9 demethylase JHDM-1 acquire longevity six to ten generations after recovery from a thaw – on average, *jhdm-1* mutant lifespan is 23.5% longer than wild-type populations of the same generation (Supplementary Fig. 1a, Supplementary Table 6). Similarly, mutants lacking WDR-5 (a core component of the H3K4 methyltransferase MLL/COMPASS complex) acquire longevity after twenty generations, living an average of 17.2% longer than generation-matched wild-type controls (Supplementary Fig. 1b, Supplementary Table 6) (Greer et al. 2010; Lee et al. 2019). Once these mutant populations have attained longevity, their descendants tend to remain long-lived, with the lifespan extension in subsequent generations reaching a plateau that fluctuates around the average (Lee et al. 2019).

Because the longevity of *wdr-5* mutants requires a proliferating germline (Greer et al. 2010), we decided to investigate the involvement of germline-to-soma signaling. The nuclear hormone receptor DAF-12 regulates developmental decisions and multiple pathways implicated in lifespan – it has also been implicated in the transgenerational longevity of *spr-5* mutants that lack a H3K4 demethylase (Greer et al. 2016). We assessed two alleles of the *daf-12* gene: the *m20* reference allele, which is a nonsense mutation, and the *rh61rh411* null mutation. Consistent with prior observations, *daf-12 (m20)* mutants never had extended lifespans (Fig. 1 and 2, Supplementary Table 1) (Larsen et al. 1995; Gems et al. 1998; McCulloch and Gems 2007). Their lifespan either resembled generation-matched wild-type controls (*P* > 0.05 across observations, log-rank test) or was shorter than wild-type populations (*P* < 0.01 across observations, log-rank test). We also did not observe a consistent change in *daf-12* mutant lifespan over generational time (Fig. 1 and 2, Supplementary Table 1).

We first tested whether *daf-12* activity was required for the establishment of longevity: could mutant populations lacking DAF-12 function ever become long-lived? We followed populations of *jhdm-1; daf-12 (m20)* double mutants across generational time and never saw longevity appear in the population, even after *jhdm-1* single mutants had become long-lived in F6 and F9 (Fig. 1c and d). The suppression of longevity was even more pronounced in *wdr-5; daf-12* double mutants, which had a shorter lifespan than wild-type controls (Fig. 1f and g) (*P* < 0.01 for both comparisons, log-rank test); we note that *wdr-5; daf-12* double mutants also had shorter lifespans than earlier generations of *wdr-5* mutant populations, even before mutant populations acquired longevity (*P* < 0.05 for all observations, log-rank test) (Fig. 1f-g). These observations were replicated using a *daf-12 (rh61rh411)* null mutation, which has a shorter lifespan than generation-matched wild-type populations (Supplementary Fig. 2) (Fisher and Lithgow 2006). Therefore, although DAF-12 has a limited effect on lifespan in otherwise wild-type populations, it was required for the establishment of transgenerational longevity in both *jhdm-1* mutants and *wdr-5* mutants.

### The maintenance of longevity has differing requirements for *daf-12* activity

For some epigenetic phenomena, the mechanisms that allow the establishment of a trait differ from those that maintain it for multiple generations (Burton and Greer 2022; Fitz-James and Cavalli 2022). Therefore, we tested whether *daf-12* activity is required to maintain longevity once it has been acquired (genetic schema shown in Fig. 2a and d). To remove *daf-12* activity only in late generations, we took advantage of a remarkable aspect of the longevity of *jhdm-1* and *wdr-5* mutants: it can be inherited by wild-type descendants for four generations, even when their descendants are genetically wild-type (Maures et al. 2011; Lee et al. 2019). To allow for the transgenerational inheritance of longevity through the maternal lineage, we crossed long-lived populations of *jhdm-1* mutants with *jhdm-1; daf-12* double mutant males to bring in the *daf-12* mutation (Fig. 2c) – we then compared the lifespan of outcrossed *jhdm-1; daf-12* double mutants to their *jhdm-1* single mutant cousins (Fig 2b). Using a similar approach, we generated *wdr-5; daf-12* double mutants with *wdr-5/+; daf-12* males (since *wdr-5* homozygous males do not mate) (Fig. 2f).

We found that outcrossed *jhdm-1; daf-12* double mutants were as long-lived as their *jhdm-1* single mutant cousins (*P* > 0.1 compared to *jhdm-1*, log-rank test): across multiple replicates, *jhdm-1; daf-12* double mutants lived 14.6% longer than generation-matched wild-type controls, and *jhdm-1* single mutants lived 16.4% longer (*P* < 0.001 compared to WT, log-rank test) (Fig. 2b, Supplemental Table 2). In contrast, *wdr-5; daf-12* double mutant populations were never long-lived: across multiple replicates, *wdr-5; daf-12* double mutant lifespan was the same as, or shorter than, wild-type controls (Fig. 2e, Supplemental Table 2). Across multiple replicates, the lifespan of *wdr-5; daf-12* double mutants was 19% shorter than that of their long-lived *wdr-5* single mutant cousins (*P* < 0.05, log rank test).

Altogether, *daf-12* activity is required for maintaining longevity in *wdr-5* mutants, but not in *jhdm-1* mutants. It is notable that this requirement differs from what we observed for establishment of longevity, in which *daf-12* activity was necessary in both mutants.

### Mutations in *daf-12* do not affect fertility in transgenerational longevity mutants

In addition to modulating lifespan, DAF-12 also regulates multiple aspects of developmental timing, including the decision to enter dauer diapause under unfavorable conditions (Cassada and Russell 1975; Riddle et al. 1981; Antebi et al. 2000). We wondered whether the genetic interaction between *daf-12* and transgenerational longevity mutants is preserved earlier in development. We first examined progeny number as a proxy for gametogenesis and embryogenesis. Consistent with previous studies, *daf-12 (m20)* single mutants produced 19.8% fewer progeny than wild-type mothers (193 progeny compared to 240 progeny, *P* < 0.05, ANOVA) (Fig. 3, Supplementary Table 3) (Antebi et al. 1998; Gerisch et al. 2001; Ow et al. 2021). Fertility was not affected in *jhdm-1* single mutants, nor in *jhdm-1; daf-12 (m20)* double mutants (*P* > 0.05 for all comparisons, ANOVA). This genetic relationship was also recapitulated with the *rh61rh411* null allele, which had previously been shown to have no fertility defect (Fisher and Lithgow 2006). As previously observed, *wdr-5* single mutants produced 66% fewer progeny than wild-type controls (*P* < 0.0001, ANOVA) (Fig. 3, Supplementary Table 3) (Simonet et al. 2007; Li and Kelly 2011; Lee et al. 2019). The fertility of *wdr-5; daf-12* double mutants resembled *wdr-5* single mutants (*P* > 0.9, ANOVA), with double mutants producing 59.2% fewer progeny than wild-type controls. The impact of a *wdr-5* mutation on fertility appeared to be genetic, rather than epigenetic, since fertility did not change over generational time (Lee et al. 2019). Overall, the loss of *daf-12* activity does not affect fertility in either of these mutant backgrounds.

**Figure 3.**
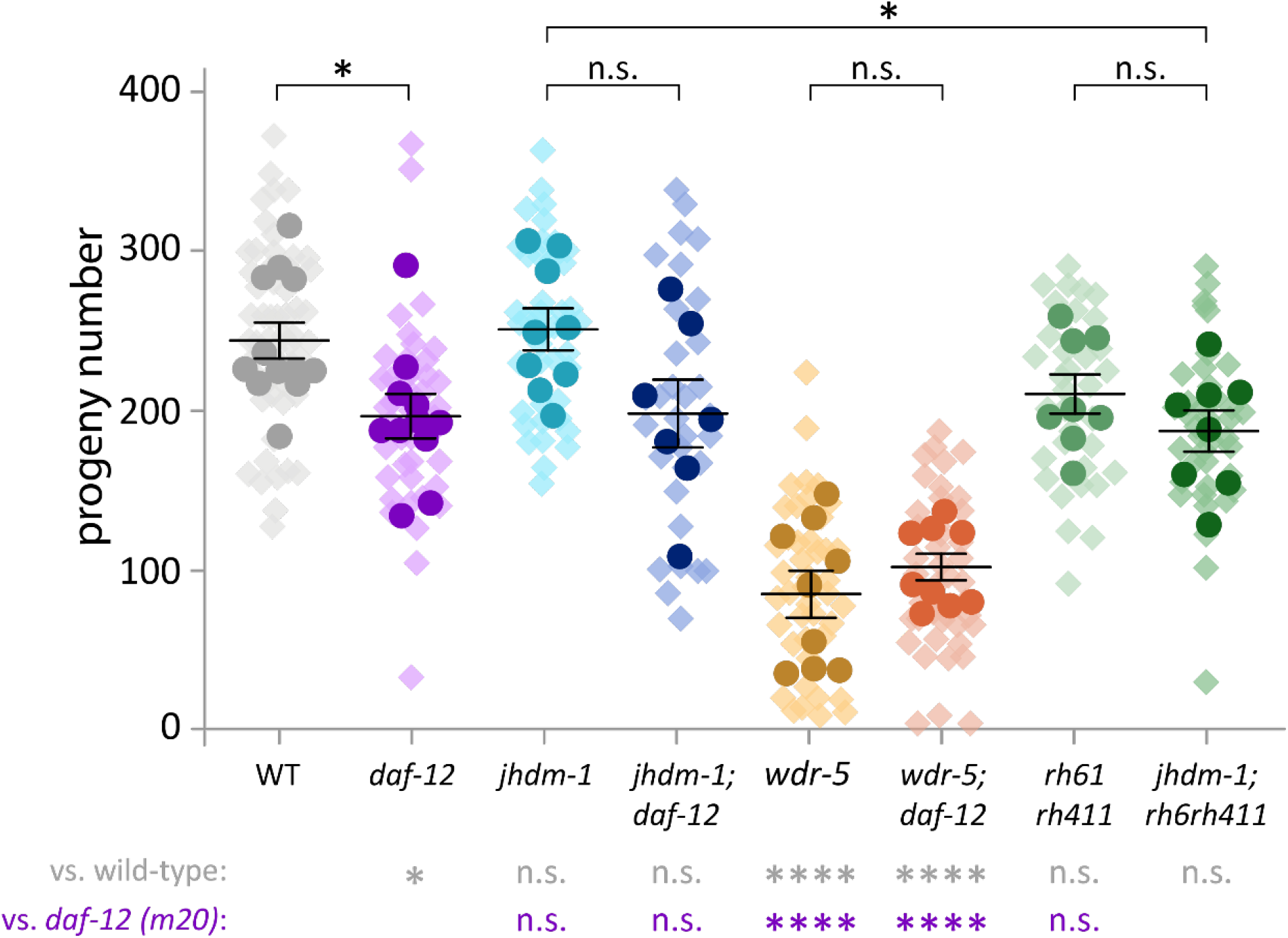
Mutations in *daf-12* do not affect fertility. Progeny number across genotypes. Darker colored circles indicate averages for each biological replicate; lighter colored diamonds indicate progeny number from individual mothers (technical replicates). The longer line represents the average progeny number across all biological replicates, with whiskers representing standard deviation. Statistical analysis was performed between biological replicates as indicated by brackets or below graph using a one-way ANOVA with a post-hoc Tukey test (n.s. = not significant, \**P* < 0.05, \*\**P* < 0.01, \*\*\*\**P* < 0.0001). Summary data and statistics for each biological replicate are shown in Supplementary Table S3.

### Mutations in *jhdm-1* and *wdr-5* enable dauer entry in a *daf-12* background

The dauer pathway is activated at the L1-to-L2 larval transition by environmental stress, including crowding, starvation, and heat (Cassada and Russell 1975; Riddle et al. 1981; Karp 2018). Dauer entry is controlled by the DAF-12 nuclear hormone receptor, which integrates insulin-like signaling (DAF-2) and TGF-ß signaling (DAF-7) pathways to regulate dauer entry, in addition to heterochronic and miRNA targets (Antebi et al. 2000; Hochbaum et al. 2011). To test the conservation of the genetic interaction during the dauer decision, we induced dauer by exposing populations to a high temperature of 27°C for the first 44 hours of development (Fig. 4, Supplementary Table 4). Consistent with prior studies, wild-type populations consisted of 22.6% dauer animals after heat shock, with variable penetrance between technical replicates, ranging from 3% to 68% dauer (Ailion and Thomas 2000; Ailion and Thomas 2003). Dauer entry in *jhdm-1* mutants or *wdr-5* mutants resembled wild-type (*P* > 0.8 for both comparisons, ANOVA). As a negative control, we assessed two alleles of the *daf-12* gene that are strongly dauer defective (daf-d) – *m20* and *rh61rh411* – and confirmed their failure to enter dauer (Fig. 4) (Antebi et al. 1998; Antebi et al. 2000; Hochbaum et al. 2011).

**Figure 4.**
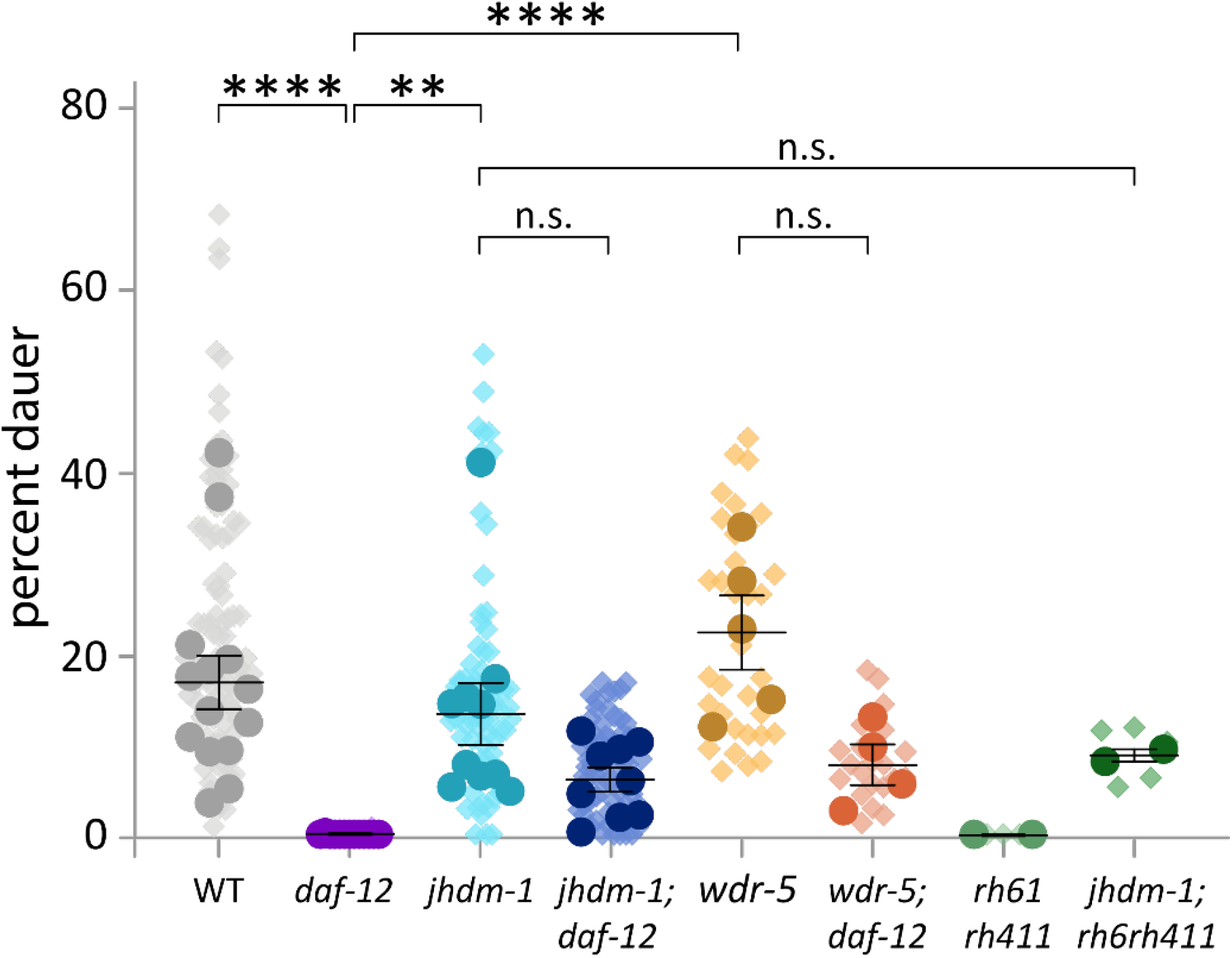
Mutations in either *jhdm-1* or *wdr-5* can suppress the dauer defect of *daf-12* mutants. The percentage of populations entering dauer under heat stress. Darker-colored circles indicate averages for each biological replicate; lighter-colored diamonds indicate the percentage for each plate scored (technical replicate). The longer line represents the average percentage dauer across all biological replicates, with whiskers representing standard deviation. Statistical analysis was performed between biological replicates using a one-way ANOVA with a post-hoc Tukey test (n.s. = not significant, \*\**P* < 0.01, \*\*\*\**P* < 0.0001). Summary data and statistics for each biological replicate are included in Supplementary Table S4.

Surprisingly, the complete absence of dauer animals in *daf-12 (m20)* mutants was rescued in both *jhdm-1; daf-12* double mutants (6% dauer) and *wdr-5; daf-12* double mutants (7.7% dauer). Although the difference between each double mutant and the *daf-12 (m20)* single mutant was not statistically significant, the presence of any dauer animals at all is striking – we never observed dauers in *daf-12 (m20)* single mutants across 14 biological replicates from two independent observers (Supplementary Table 4). Because the *daf-12 (m20)* allele is not a null, we replicated this observation with *jhdm-1; daf-12 (rh61rh411)* double mutants (8.7% dauer). However, unlike lifespan, dauer entry does not change across generational time: there was no significant difference in dauer animals between normal-lived and long-lived populations (*P* > 0.2 for all genotypes, t-test) (Supp. Fig. 3).

We verified the presence of dauer animals in double mutant populations by scoring resistance to 1% SDS. Dauer animals have a thicker cuticle that allows them to survive in SDS detergent for hours, whereas wild-type animals die within a few minutes (Cassada and Russell 1975; Karp 2018). After 30 minutes of SDS exposure, we observed living animals in both *jhdm-1; daf-12* double mutant and *wdr-5; daf-12* double mutant populations, but never in *daf-12 (m20)* or *daf-12 (rh61rh411)* single mutants (data not shown). Overall, mutations in either *jhdm-1* or *wdr-5* rescued the dauer defect of *daf-12* mutants, indicating that changes in chromatin landscapes may affect DAF-12 downstream signaling at an early developmental timepoint.

### *wdr-5* mutation in *daf-12* background triggers developmental delay

The impact on dauer entry led us to test this genetic interaction in the context of an unchallenged development and favorable conditions. In wild-type populations, 99% of wild-type animals were L2 larvae after 24 hours; by 48 hours, they were L4 larvae; and at 72 hours, nearly all animals had reached adulthood (Fig. 5, Supplementary Table 5). Mutations in *daf-12* or *jhdm-1* do not affect the timing of larval development, and as expected, *jhdm-1; daf-12* double mutants also strongly resemble wild-type controls (*P* > 0.1 for all comparisons, Fisher’s exact test) (Fig. 5A). In contrast, *wdr-5* single mutants experienced a slight delay compared to age-matched wild-type controls (*P* < 0.0001 for all timepoints, Fisher’s exact test) (Fig. 5B) – at 72 hours, when nearly all wild-type animals are adults, 20% of *wdr-5* mutants were still L4 stage or younger. Similar to what we observed with fertility and dauer entry, developmental timing was not affected by generational time in either *jhdm-1* or *wdr-5* single mutants (Supplementary Table 5).

**Figure 5.**
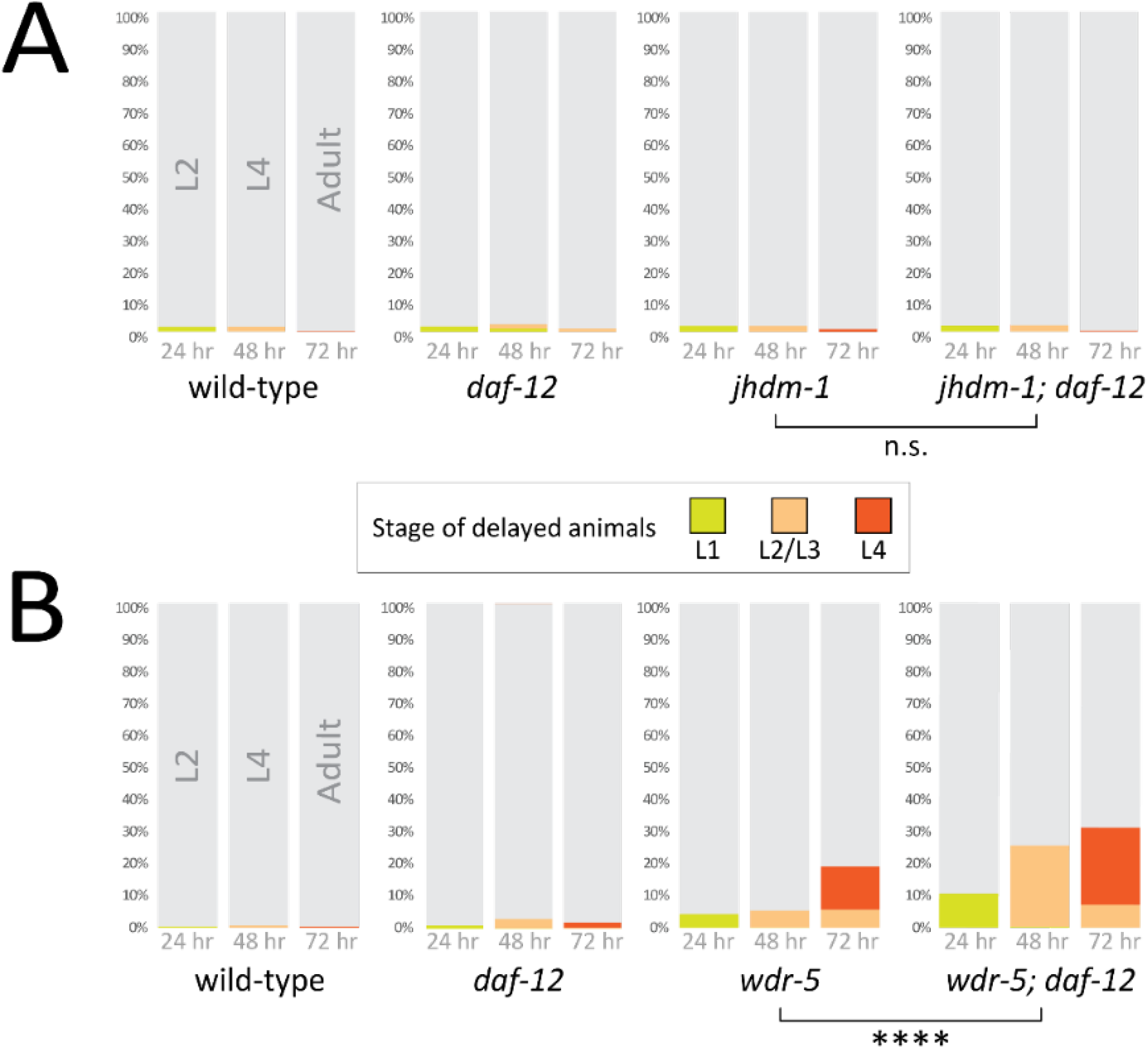
Mutations in *daf-12* enhance developmental delay in *wdr-5* mutants. Stacked bar graphs indicating the percentage of each population at the specified stage in *jhdm-1; daf-12* double mutants (**A**) and *wdr-5; daf-12* double mutants (**B**). Gray bars represent the stage of wild-type populations at 24 hours (L2 larvae), 48 hours (L4 larvae), and 72 hours (adults); colored bars represent younger larval stages as indicated in the legend. Statistical analysis was performed using a Fisher’s exact test comparing the single mutants to *daf-12* double mutants at the same timepoint; significance was the same across each of the three timepoints (n.s. = not significant, \*\*\*\**P* < 0.0001). Additional statistical comparisons, summary statistics, and statistics for each biological replicate are included in Supplementary Table S5.

The developmental delay of *wdr-5* single mutants became more pronounced in *wdr-5; daf-12* double mutants (*P* < 0.0001 for all timepoints compared to wild-type controls and *wdr-5* single mutants, Fisher’s exact test) (Fig. 5B, Supplementary Table 5). At 72 hours, 35% of the double mutants had yet to reach adulthood, with some animals still at the L2 larval stage. Both *wdr-5* single mutants and *wdr-5; daf-12* double mutants experienced some delay in hatching: at 24 hours, the single mutant had a 4% delay and the double mutant had a 12% delay) – this delay suggested defects during embryogenesis, which was consistent with a higher embryonic lethality previously reported for *wdr-5* single mutants (Li and Kelly 2011; Lee et al. 2019). However, the embryonic delay could not account for the full delay observed at 72 hours, which suggested an additional defect with larval development. We note that the presence of larvae in populations at this timepoint represented a delay, rather than an arrest, as they eventually become adults after 96 hours (data not shown). We also noticed that *wdr-5; daf-12* double mutants had a higher incidence of defective vulva development, including the multi-vulva (Muv) and protruding vulva (pVulv) phenotypes, which were never observed in either single mutant population (data not shown). Because the delayed development of *wdr-5* single mutants was exacerbated by a *daf-12* mutation (*P* < 0.0001 for all timepoints compared to *wdr-5* controls, Fisher’s exact test), changes in MLL/COMPASS function may affect DAF-12 signaling during developmental decision-making. Future studies will examine how aberrant chromatin landscapes affect earlier embryonic development.

## DISCUSSION

In this study, we explore the relationship between development and aging by testing the genetic interaction between the DAF-12 signaling pathway and heritable chromatin landscapes. We find that each influences the other at different points during the *C. elegans* life cycle: during development, *daf-12* is required for normal progression; during dauer diapause, chromatin landscapes bypass the need for DAF-12 signaling; and during aging, *daf-12* activity is necessary for heritable heterochromatin to confer longevity. Taken together, these results highlight how early developmental gene programs may be coopted by heritable chromatin landscapes to influence an individual’s biology throughout its life cycle.

Organismal development requires the precise spatial and temporal coordination of biological pathways with environmental cues – therefore, developmental gene networks have evolved to be robust in the face of highly variable inputs. However, a growing body of work suggests that these programs can continue to impact an individual’s physiology throughout its life, well beyond the constraints of original selection pressures during development (Silva-García 2023; Raffington 2024). Ever since researchers discovered that aging is controlled genetically (Klass 1983; Friedman and Johnson 1988; Kenyon et al. 1993), the field has raced to identify the specific pathways involved (Kenyon 2010; Tissenbaum 2015). One recent overarching theory of aging is the programmatic theory, which attempts to resolve how evolutionary mechanisms allow for the persistence of alleles that become deleterious as individuals age: developmental gene regulatory networks are mechanistically programmed to allow for success early in life, but can become detrimental to health when they remain active in aged individuals (De Magalhães and Church 2005; Blagosklonny 2006; Gems 2022).

The programmatic theory elegantly synthesizes two earlier hypotheses for why aging occurs. The first of these theories, the disposable soma hypothesis, is based on the high cost of repairing molecular damage – with limited resources, organisms must navigate a trade-off between self-maintenance or investing in the next generation. Since the soma only needs to be maintained until reproduction, organisms prioritize repair in germline cells, leaving somatic cells to accumulate damage and therefore age (Kirkwood 1977; Kirkwood and Holliday 1979; Mukhopadhyay and Tissenbaum 2007). The second theory, the antagonistic pleiotropy hypothesis, proposes that alleles that benefit young individuals can have detrimental effects on aged individuals (Williams 1957). Because these alleles provide a fitness benefit, they are under positive selection and must be considered wild type – therefore, aging-related diseases are caused by the activity of a wild-type genome. This theory is supported by empirical work demonstrating a fitness cost when lifespan is genetically extended (Jenkins et al. 2004). Bringing together these hypotheses, the programmatic theory is grounded in the principle that natural selection acts upon traits that increase fitness. Because events that occur earlier in life have a greater impact on reproductive fitness, selection has primarily refined developmental programs that work well earlier in life. Aging and aging-related diseases are simply the unintended consequences of having these same programs continue into adulthood. This theory also helps to explain one notable feature of aging: its variability between individuals of the same species. Even completely isogenic populations of *C. elegans* reared in the same dish will experience lifespans that can vary widely, from as short as 10 days to as long as 30 days. Developmental programs must be precise and stereotyped early in life, but stochasticity develops as precision is loosened during adulthood (Gems 2022).

As biological processes, both aging and development share the need to keep track of time. The fact that these processes exist on the same continuum of life history has been supported by the discovery of DNA methylation clocks among mammals and other vertebrates: they tick rapidly during development and continue ticking (albeit more slowly) through adulthood (Horvath and Horvath 2013; Jones et al. 2015; Raj and Horvath 2020). Two observations provide further evidence that aging and development share common mechanisms. First, many epigenetic clock sites are also associated with developmental programs, including the Polycomb regulation of Hox genes (De Magalhães 2012; Raj and Horvath 2020). Second, conserved sites of DNA methylation can predict animal age across a wide variety of mammalian taxa, indicating that these loci reflect an underlying shared gene regulation (Lu et al. 2023). The fact that clock changes can be detected in young, developing individuals further bolsters the idea that development and aging lie on a unified continuum.

The connection between development and aging is highlighted by the common role of the DAF-12 nuclear hormone receptor. DAF-12 was first discovered in genetic screens for components regulating dauer diapause, a key developmental timepoint (Riddle and Albert 1997; Antebi et al. 2000). It was then shown to act as an important bridge between environmental cues and the heterochronic pathway through the regulation of microRNAs (Antebi et al. 2000; Hammell et al.). Later, DAF-12 was shown to be necessary for lifespan extension in animals with ablated germ cells (Hsin and Kenyon 1999). DAF-12 controls target gene expression in two ways: by controlling the nuclear localization of the DAF-16/forkhead (FOXO) transcription factor (Hsin and Kenyon 1999; Berman and Kenyon 2006; Mukhopadhyay et al. 2006),x0 or by binding DNA directly (Hochbaum et al. 2011). When the germline is ablated in *C. elegans*, DAF-16/FOXO enters the nucleus only in intestinal cells – *daf-12* gene activity is necessary for both the lifespan extension and DAF-16 nuclear localization, highlighting its role in germline-to-soma communication (Arantes-Oliveira et al. 2002; Berman and Kenyon 2006; Kenyon 2010).

Each of the transgenerational longevity mutants used in this study, *wdr-5* and *jhdm-1*, acquire longevity in generations with higher levels of heterochromatin (Lee et al. 2019). We previously proposed the following model for heterochromatin accumulation. In wild-type animals, COMPASS deposits active H3K4me at actively transcribed germline genes. During epigenetic reprogramming, the methyltransferase MET-2 deposits repressive H3K9me2 in genomic loci that lack H3K4me (Kerr et al. 2014; Carpenter et al. 2021; Delaney et al. 2022). However, during epigenetic reprogramming in COMPASS mutants, more repressive H3K9me2 is deposited in the absence of active H3K4me. With each successive generation, genomic levels of H3K9me2 continue to increase, which ultimately causes longevity in the population.

To understand how the accumulation of heterochromatin in the germline could influence somatic aging, we tested the requirement for DAF-12 germline-to-soma signaling. The transgenerational longevity of both *wdr-5* mutants and *jhdm-1* mutants shares an underlying cause, so we expected that both mutants would have similar relationships with DAF-12. Although this was true for the *establishment* of longevity (*daf-12* gene activity was necessary for both mutant populations to become long-lived), it was not the case for *maintaining* longevity once it has been acquired: removing the activity of *daf-12* had no effect on long-lived *jhdm-1* mutant populations, although it did abrogate longevity in *wdr-5* mutants. The difference in epistatic relationships highlights two points. First, that the transgenerational acquisition of longevity in both mutant backgrounds shares a common mechanism (the accumulation of repressive H3K9me2), and that this mechanism requires DAF-12, perhaps because of its role in germline-to-soma signaling. Second, the downstream impacts of inherited H3K9me2 must affect somatic phenotypes via distinct mechanisms, including whether DAF-12 is involved in larval development or lifespan extension after longevity has been acquired by the population. Current projects are examining how gene expression and chromatin accessibility are affected in these mutants, which will provide insight on how they intersect with DAF-12 regulation and other longevity pathways.

We also noted that the loss of *daf-12* activity differently affected lifespan in these genetic backgrounds. Although lifespan in *wdr-5* single mutants was occasionally shorter than wild-type controls, it was rarely statistically significant. However, *wdr-5; daf-12* double mutants frequently had significantly shorter lifespans than wild-type controls, and often shorter than *wdr-5* single mutant controls. When probing this genetic relationship earlier in development, we also saw a similar difference: *wdr-5; daf-12* double mutants experienced a more severe delay than *wdr-5* single mutants, whereas *jhdm-1; daf-12* double mutants developed normally. The combination of these phenotypes highlights their severity: because we start our lifespan assays using a synchronized laying period, one might expect that the delayed development of *wdr-5; daf-12* double mutants gives them a slight boost – by reaching adulthood 1-2 days later than control populations, they might be expected to live slightly longer. However, their shorter lifespans indicate that problems arising during development persist and affect health during adulthood.

We were surprised to find that mutations in *wdr-5* or *jhdm-1* could suppress the strong dauer-defective phenotype of both *daf-12* alleles. Dauer entry is controlled through two parallel pathways that converge on the DAF-12 nuclear hormone receptor: the DAF-7/TGF-β pathway detects dauer pheromone, whereas the DAF-2/insulin-like signaling pathway detects the presence of food (Wolkow and Hall 2015). In the presence of dauer pheromone, DAF-12 is prevented from binding its ligand, and instead binds the co-repressor DIN-1S to repress reproductive development and promote dauer entry (Ludewig et al. 2004; Mahanti et al. 2014). The *daf-12 (m20)* allele used throughout this study is hypomorphic due to an early stop codon in the DNA binding domain that truncates three out of four isoforms of the *daf-12* transcript. We also repeated some experiments using the *rh61rh411* allele, which is a putative null due to mutations in both the DNA-binding and ligand-binding domains, one of which is an early stop codon that truncates all four isoforms (Antebi et al. 2000; Hochbaum et al. 2011). It is possible that the widespread changes in chromatin landscapes caused by the generational loss of *jhdm-1* or *wdr-5* allow truncated forms of the DAF-12 receptor to access DNA and trigger the dauer developmental program. Alternatively, the altered chromatin landscape may somehow create a permissive state for dauer entry in response to stress, bypassing the need for DAF-12 transcriptional regulation entirely. Finally, the interactions between DAF-12 and lipid metabolism may shed light on longevity in these mutant backgrounds. COMPASS mutants accumulate more fat, with an enrichment of mono-unsaturated fatty acids, which are produced by a pathway that includes *fat-7* (Han et al. 2017). A recent preprint has shown that oleic acid, the lipid metabolite produced by *fat-7*, modifies the activity of DAF-12 to promote lifespan as well as fertility (Nichitean et al. 2023).

Across metazoans, nuclear hormone receptors direct complex transcriptional programs in response to multiple inputs (Gustafsson 2016; Weikum et al. 2018). As the downstream transcriptional effectors of signaling pathways, nuclear hormone receptors have the remarkable ability to act as either an activator or repressor, depending on the specific context in which they are deployed – transcriptional impact can be modified by minor changes to receptor structure caused by ligand binding or association with co-factors, which may themselves vary in availability depending on the specific developmental timepoint (Hochbaum et al. 2011; Taubert et al. 2011; Hazarika et al. 2024). DAF-12 is homologous to human metabolic sensors like vitamin D receptor and liver X receptor, and many components of the endocrine signaling network are conserved between *C. elegans* and mammals (Fielenbach and Antebi 2008). For liver X receptor isoforms, molecular dynamic simulations have demonstrated that allosteric interactions can enact significant differences in function (including transcriptional regulation) despite high structural similarity (Choudhuri and Okafor 2026). It is not clear how local chromatin states impact DAF-12 binding in its ligand-bound or unliganded state, let alone how DNA binding may be affected by its formation as a homodimer or heterodimer. The variable transcriptional output from a single structure for homologous isoforms may provide hints at why altered heterochromatin landscapes in *jhdm-1* and *wdr-5* mutants bypasses ability of *daf-12* mutants to enter dauer. Future work will leverage molecular dynamic simulations to model how different *daf-12* alleles affect isoform structure and ability to bind in different chromatin contexts.

Nuclear hormone receptors like DAF-12 play a key role in integrating environmental cues with endocrine signaling to enact key developmental decisions. DAF-12’s role in promoting communication from the germline to the soma highlights how this integration also requires significant communication between tissue types: one tissue or organ can sense changes in the local environment and initiate signals to other tissues. In *C. elegans*, a common theme for this kind of systemic activation has been through neurons, which have the ability to sense a wide range of cues and initiate trans-cellular signaling (Durieux et al. 2011; Posner et al. 2019; Smith et al. 2020). In the related nematode species, *Pristionchus pacificus*, the same DAF-12 signaling pathways regulate both dauer entry and olfactory chemotaxis by controlling neuronal fate (Carstensen et al. 2021). One of the more intriguing consequences of signaling between neurons and other tissues has been to the germline, which provides a mechanism for epigenetic inheritance of perception or behavior by subsequent generations. For example, exposure to pathogenic bacteria generates intergenerational and transgenerational avoidance for up to four generations (Kaletsky et al. 2020; Sengupta et al. 2024; Pender et al. 2025). Future studies will investigate how the transgenerational inheritance of heterochromatin impacts neuronal function and germline communication.

Overall, we have shown that the transgenerational acquisition of longevity in *jhdm-1* and *wdr-5* mutants co-opts an existing developmental pathway mediated by DAF-12 signaling. The genetic interactions between the transgenerational mutations and *daf-12* mutations indicate that chromatin regulation has a critical function in developmental transitions across an individual’s life history.

## DATA AVAILABILITY

Strains are available from the *Caenorhabditis* Genetics Center or upon request. The authors affirm that all data necessary for confirming the conclusions of the article are present within the article, figures, and tables.

## ACKNOWLEDGEMENTS

We would like to thank Eric Greer, Swathi Arur, Sarah Hall, David Katz, and Michelle Mondoux for providing suggestions and insight for this project. Strains were provided by the *Caenorhabditis* Genetics Center (RRID:SCR_007341), which is funded by the National Institutes of Health (NIH) through the Office of Research Infrastructure Programs (P40 OD010440).

---

**Figure S1.**
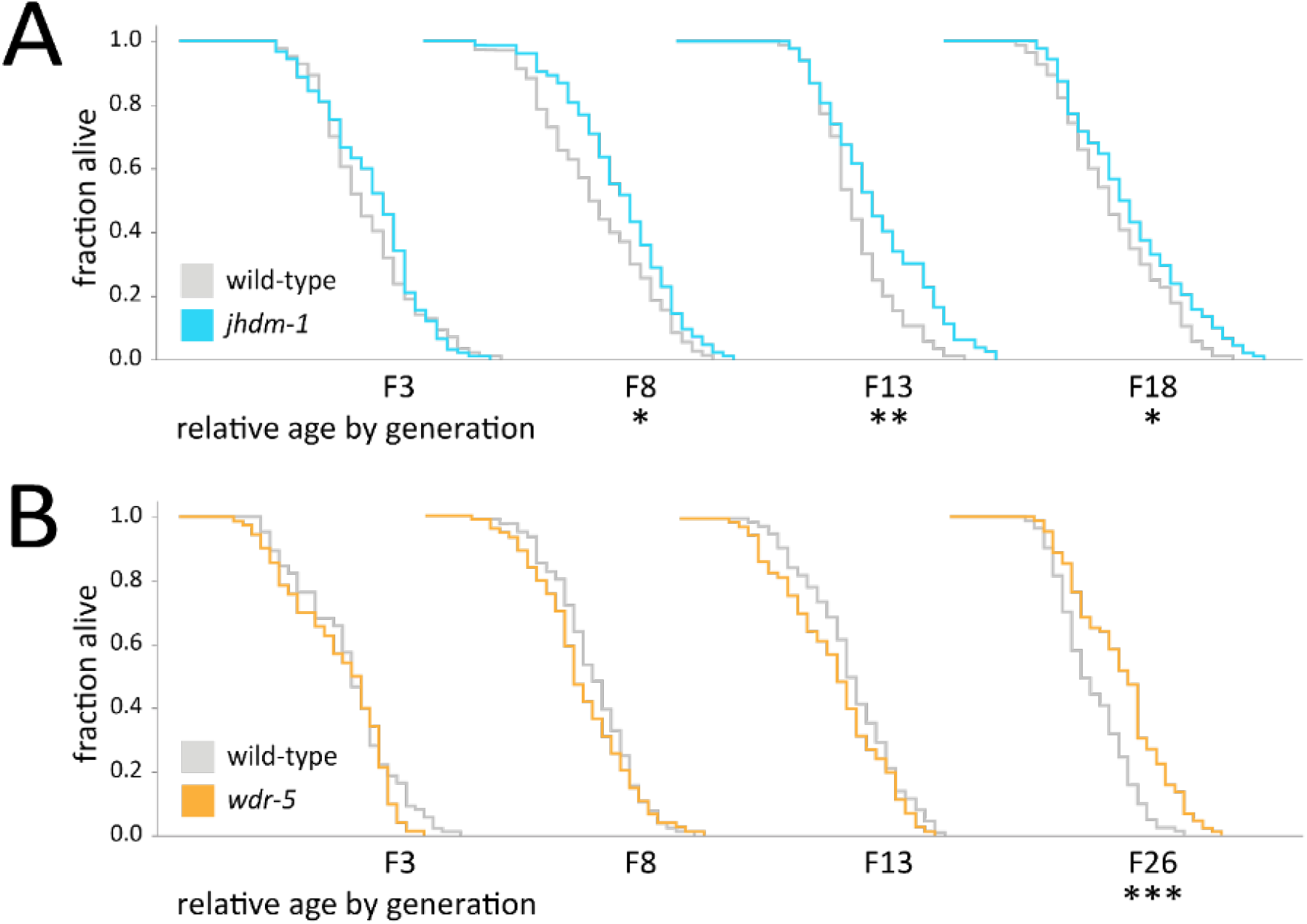
*jhdm-1* and *wdr-5* mutants acquire longevity across many generations. (**A**, **B**) Analysis of relative lifespan in *jhdm-1* mutants (**A**, blue) and *wdr-5* mutants (**B**, orange) compared to generation-matched wild-type populations (gray). P0 populations were descended from the L1 larvae recovered from a thaw. For each generation, the x-axis represents 35 days. Statistical analysis was performed using a log-rank test compared to generation-matched wild-type populations (\**P* < 0.05, \*\**P* < 0.01, \*\*\**P* < 0.001). Median lifespan, statistics, and additional replicates are shown in Supplementary Table S6.

**Figure S2.**
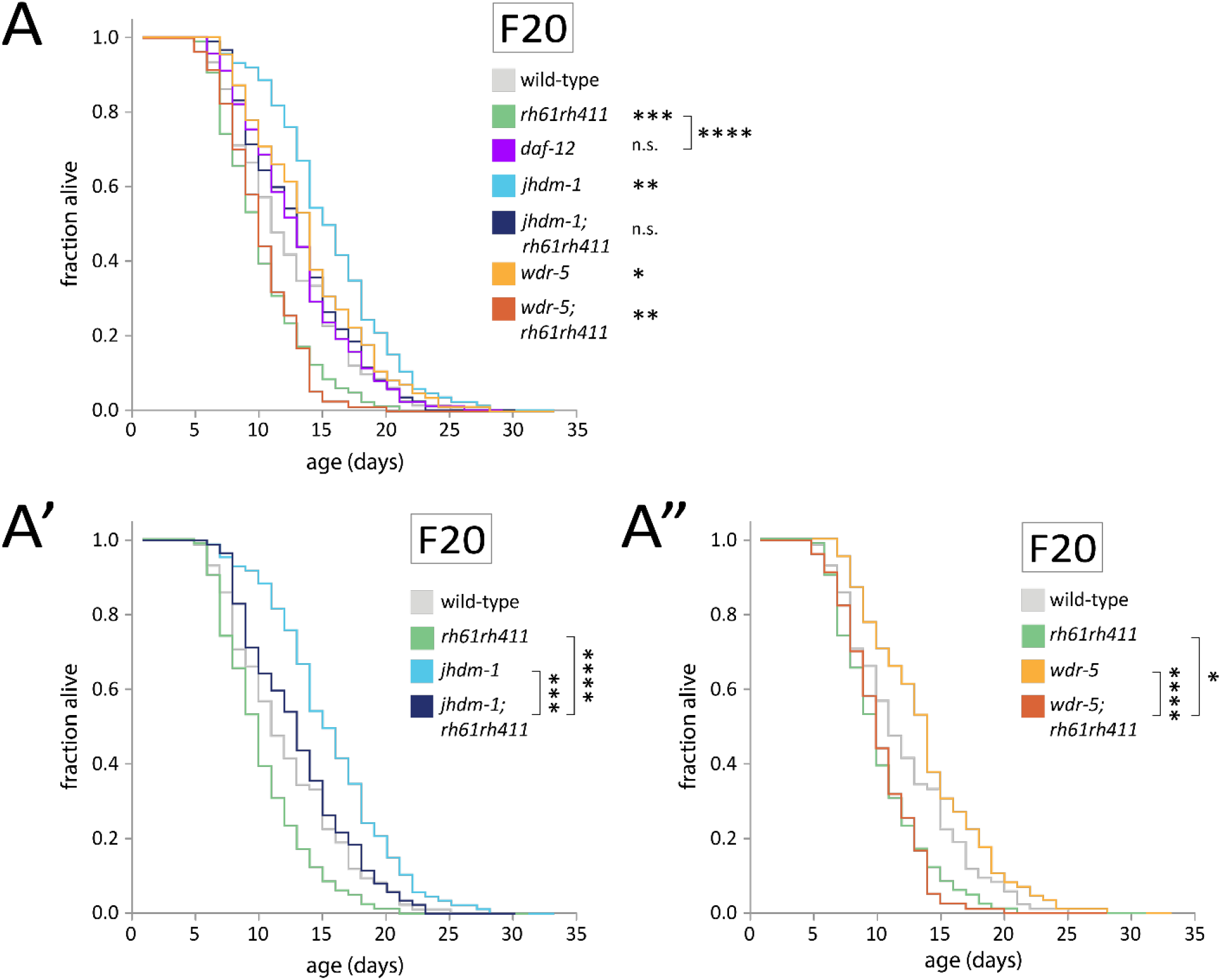
*daf-12 (rh61rh411)* also suppresses the acquisition of longevity. (**A**) Lifespan in transgenerational *jhdm-1* mutant (light blue) and *wdr-5* mutant (yellow) populations, including *daf-12 (m20)* single mutants (purple) compared to *daf-12 (rh61rh411)* single mutants. Data are also shown separated into *jhdm-1; daf-12 (rh61rh411)* double mutants (dark blue) (**A’**) and *wdr-5; daf-12 (rh61rh411)* double mutants (dark orange) (**A”**). Statistical analysis was performed using a log-rank test compared to generated-matched wild-type populations (shown next to genotype) or between genotypes (indicated by bracket) (\**P* < 0.05, \*\**P* < 0.01, \*\*\**P* < 0.001, \*\*\*\**P* < 0.0001, n.s. = not significant). Median lifespan and statistics are included in Supplementary Table S6.

**Figure S3.**
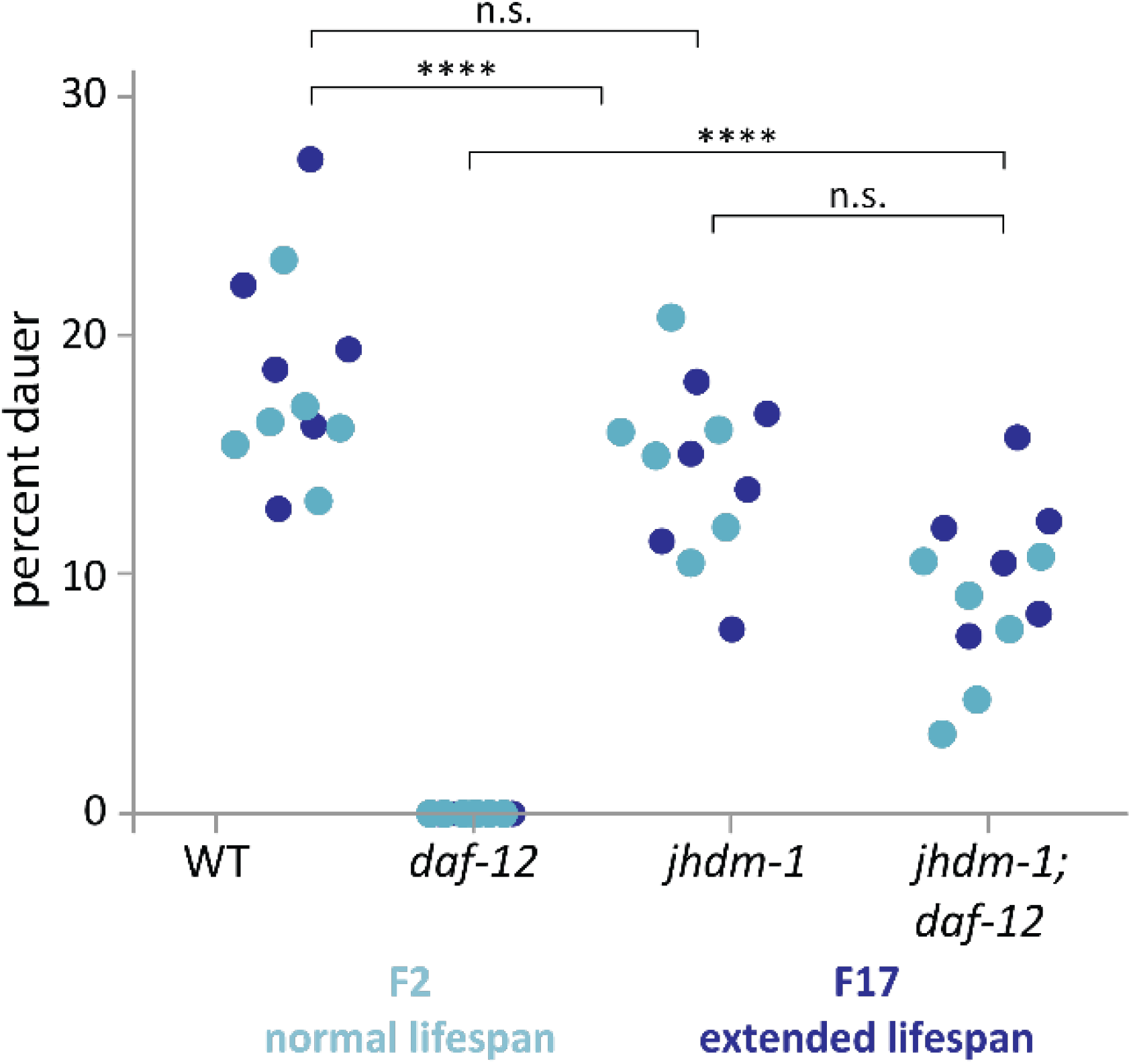
Dauer formation is not affected by generational time. The percentage of dauer entry in populations with normal lifespans (light blue) or extended lifespans (dark blue). For each genotype, comparisons between generations were performed using a t-test (n.s. = not significant, \*\*\*\**P* < 0.0001).

